# Genetic prevalence and clinical relevance of canine Mendelian disease variants in over one million dogs

**DOI:** 10.1101/2022.06.12.495799

**Authors:** Jonas Donner, Jamie Freyer, Stephen Davison, Heidi Anderson, Matthew Blades, Leena Honkanen, Laura Inman, Casey A. Brookhart-Knox, Annette Louviere, Oliver P Forman, Rebecca Chodroff Foran

## Abstract

Hundreds of genetic variants linked to Mendelian disease have been characterized in dogs to date, and commercial screening is being offered for most of them worldwide. There typically remains a paucity of information regarding the broader population frequency of newly discovered variants, as well as uncertainty regarding their functional and clinical impact on additional genomic ancestry backgrounds beyond the discovery breed. Panel screening of disease variants, commercially offered as direct-to-consumer genetic testing, provides an opportunity to establish large-scale cohorts with both genotype and phenotype data available to address open questions related to variant prevalence and relevance. In this study, we screened the largest canine cohort examined in a single study to date (1,054,293 representative dogs from our existing cohort of more than three million dogs; a total of 811,628 mixed breed dogs and 242,665 purebreds from more than 150 countries and territories) for 250 genetic disease-associated variants to understand their prevalence and distribution in the general population. Electronic medical records from veterinary clinics were available for 43.5% of the genotyped dogs, enabling follow up on the clinical impact of variants. We provide detailed frequencies for all tested variants across breeds and find that 57% of dogs carry at least one copy of a studied Mendelian disease-linked variant. We provide evidence of full penetrance for 10 variants, and at minimum plausible evidence for the clinical significance of 22 variants, on a wide variety of breed backgrounds. We further show that a reduction in genome-wide heterozygosity is associated with an increased Mendelian disease load and assess genome-wide heterozygosity levels in over 100 breeds. The accumulated knowledge represents a resource to guide discussions on disease variant presence and genetic test relevance by breed.

## Introduction

In recent years dogs have established themselves as a geneticist’s best friend by enabling biomedical research discoveries that improve our understanding of the molecular background of inherited diseases and their treatments (1, 2). Such discoveries can lead to advancements that benefit human health, while serving the dog breeder and veterinary communities through the availability of genetic tests that can help to guide breeding selections and veterinary care. With more than 300 canine Mendelian disease and trait variants identified to date (3), many of which form the basis of commercially available genetic tests, it has become more and more challenging for breeders, breed health advisors, kennel clubs and breed registries, veterinary clinicians, and scientists to stay informed about which genetic tests are relevant for which breeds and determine the level of population, functional, and clinical evidence supporting the use of each test. There are commendable initiatives to compile and make information available on inherited diseases and traits, genetic variants, genetic test availability, and genetic test relevance in dogs (e.g., (3–5)). However, one constraint of the currently available resources is the limited availability of follow-up, or routinely updated information on the frequency of genetic variants compiled after the original research discovery publication. Other current knowledge gaps related to canine disease variants stem from discovery studies published without conclusive functional evidence supporting candidate variant causality, and from the limited opportunities of original studies to explore variant impact on a broad variety of breed backgrounds beyond the initial discovery breed.

Multiplex or panel screening of known disease variants has emerged as a technologically feasible, cost-efficient, and scientifically justified approach to simultaneously screen any individual dog for a large number of disease variants (6, 7). For the first time ever, multiplex screening has recently been introduced on a large scale into the veterinary clinical setting through the inclusion of routine DNA testing in pet care plans and electronic medical records (EMRs) across more than a thousand clinics in the United States (8). Together with large-scale health surveys, this development allows for the establishment of cohorts with deep phenotypic and genetic information that hold great potential for addressing some of the knowledge gaps related to not only the genetic prevalence and distribution of disease variants in the population, but also to the clinical relevance of specific genetic testing results.

Genetic test results for Mendelian disease variants should also be considered in the broader context of population genetic diversity and the viability of the gene pool. One mechanism in which inbreeding depression (reduced survival and fitness in the offspring of close relatives) can act is through the accumulation of deleterious recessive mutations that negatively impact the health and fertility/fecundity of the individual (9). The link between genomic inbreeding levels and homozygosity for known disease variants has so far not been exhaustively explored across dog breeds with direct simultaneous genetic measurement of both variables in the same individual, although considerable overlap between regions of homozygosity (ROH) and “at risk” genotypes at 29 recessive disease loci was previously reported (10). Molecular estimates of genomic inbreeding levels for breed comparisons can be obtained through genotyping single nucleotide polymorphisms (SNPs), for example through array-based approaches, enabling simultaneous screening for genetic diseases, while providing a more accurate representation of genetic population measurements of diversity compared to pedigree-based metrics (11).

We have previously explored canine genetic disease prevalence through analyses of 152 Mendelian disease variants in around 100,000 dogs (7). In the current study, we expanded our work by genotyping a separate cohort consisting of 1,054,293 dogs for 250 genetic variants previously associated with canine disease to further understand their frequency and distribution in the general canine population. Moreover, we sought to obtain additional evidence supporting the association between the variants and the respective phenotypes to which they have previously been linked using electronic medical records (EMRs) and pet owner interviews. We additionally explored the relationship between genome-wide inbreeding level and Mendelian disease variant load.

## Results

### Quantitative analysis highlights the ubiquitous nature of canine disease variants

Of 250 genetic variants largely following Mendelian inheritance patterns and selected for screening based on their previously implicated involvement in canine inherited disorders, 207 (82.8%) were observed in at least one dog in the study population of 1,054,293 dogs. The majority of disease variants (N = 182; 87.9% of all observed disease variants) were encountered in both mixed breed and purebred dogs while 19 diseases (9.2%) were exclusive to the mixed breed population and 6 (2.9%) to the purebred population (Figure 1A; Supplementary Table 1). The maximum number of screened variants present in any one dog (two mixed breed dogs) was eight. Restricting the investigation to the subset of 242 variants genotyped in the full cohort, we observed that 57.2% of all dogs carried at least one of the tested disease variants in either hetero- or homozygous state. (Figure 1B).

**Figure 1.**
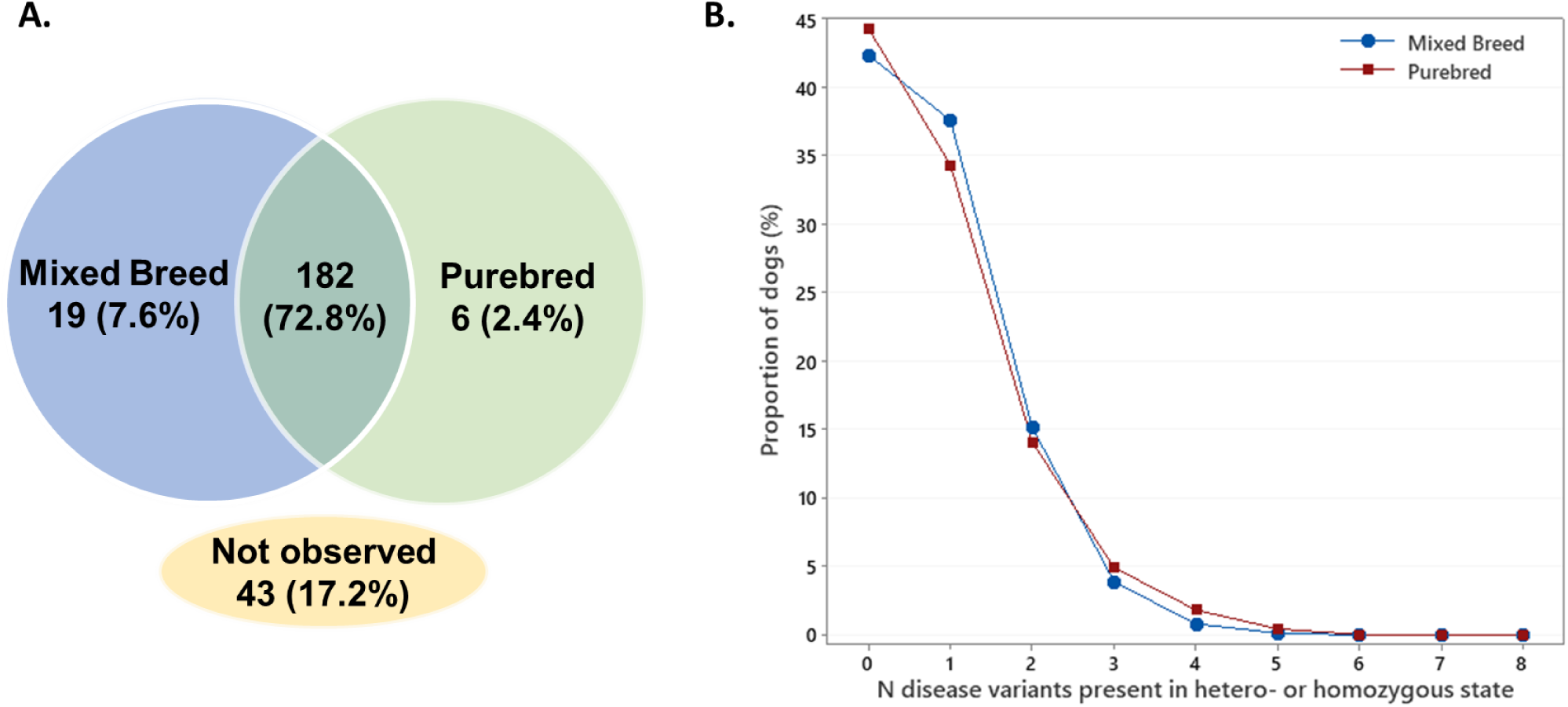
A) Presence of 250 Mendelian disease variants and B) distribution of 242 Mendelian disease variants in a cohort of 811,628 mixed breed and 242,665 purebred dogs.

### Decrease in genome-wide heterozygosity is associated with an increased Mendelian disease variant load

We have previously shown that a higher proportion of mixed breed dogs will be heterozygous for at least one of nine examined recessive disease variants compared with purebred dogs, while a higher proportion of purebred dogs will be homozygous for at least one of the same recessive disease variants compared with mixed-breed dogs (7). In the present study, we sought to further refine understanding of this pattern by leveraging the availability of genome-wide SNP-based heterozygosity estimates for individual dogs of the studied cohort, acquired as a part of commercial genetic testing. As expected, we found that purebred dogs (N = 242,665) had a lower median heterozygosity level than mixed breed dogs (N = 811,628), indicating a higher median level of genetic inbreeding in purebreds (35.98% vs 44.87% heterozygous genotyped SNP loci respectively; Mann-Whitney W = 5.03*10^11^, P < 10^-16^), and we considered these two groups separately in subsequent analyses. Figure 2 further details heterozygosity levels by breed for the 104 breed groups that were represented by ≥100 individuals while full statistics for all breeds are available in Supplementary Table 2.

**Figure 2.**
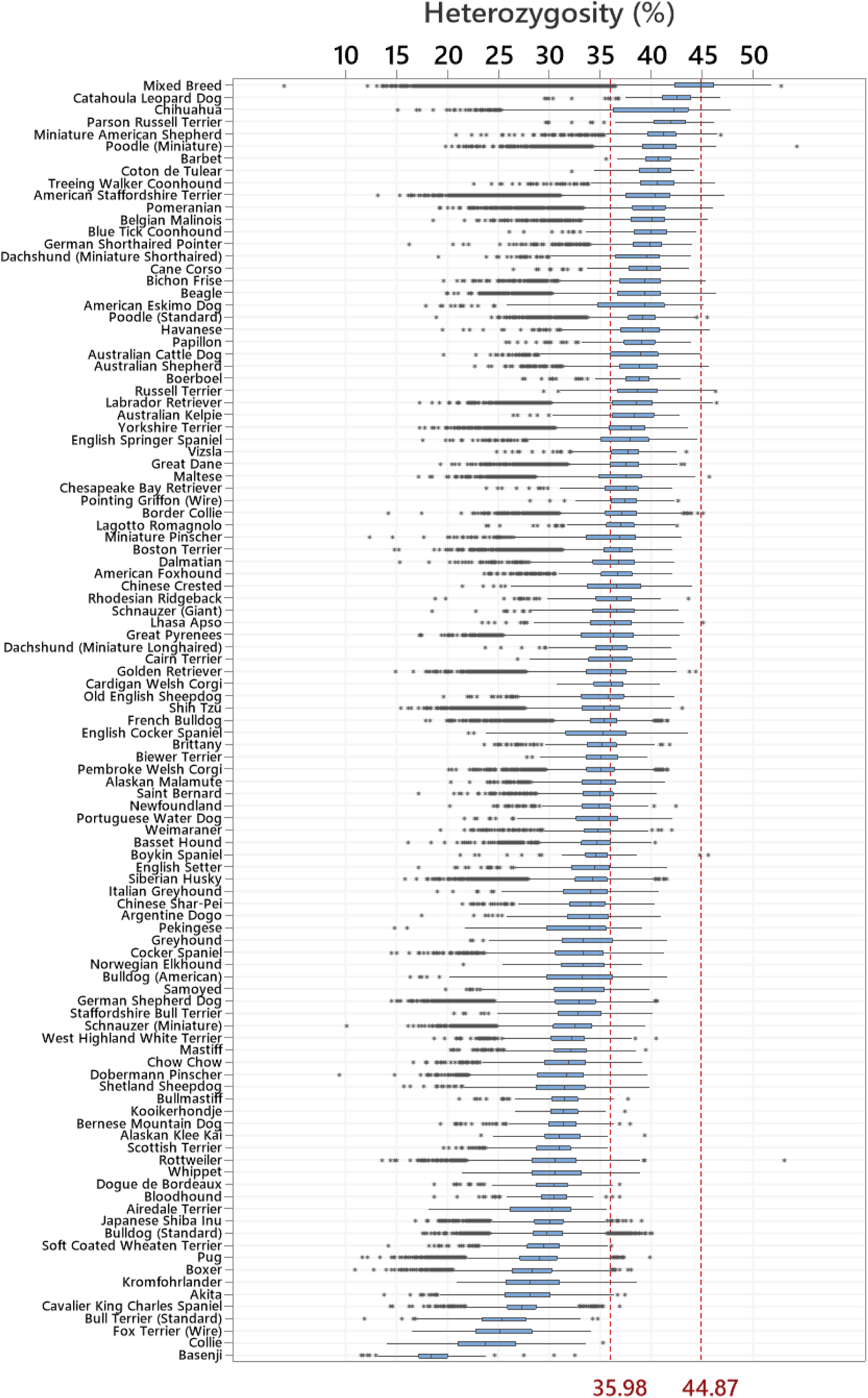
Genome-wide heterozygosity in mixed breed dogs and 104 breeds (N ≥100 dogs tested) ranked from most to least diverse. The figure shows medians and interquartile range boxes representing the middle 50% of the data, with whiskers representing the ranges for the bottom and top 25% of the data values excluding outliers (data points >1.5x the interquartile range from the boxes; shown as asterisks). The red vertical reference lines represent the median heterozygosity in the combined purebred dog group (35.98%) and mixed breed dogs (44.87%).

We examined the correlation between genome-wide levels of genetic diversity and genotypes for autosomal disease variants (178 variants following recessive and 13 semi-dominant or dominant modes of inheritance; Supplementary Table 1) present and genotyped in the full study sample. We found that increased genome-wide genetic heterozygosity was weakly correlated with the presence of an increased number of autosomal disease variants in heterozygous state in both mixed and purebred dogs (Spearman r = 0.082 and 0.082, respectively, P < 0.001; Figure 3A). Conversely, and more notably, we found that a decrease in heterozygosity manifests as an increased number of homozygous autosomal disease variants (Figure 3B). This pattern was observed in both dogs classified as mixed breed (r = -0.16, P < 0.001) and purebred (r = -0.136, P < 0.001), and persisted when the analysis was repeated including only the recessive disease variants (mixed breed r = -0.118, P < 0.001 and purebreds r= -0.155, P < 0.0001).

**Figure 3.**
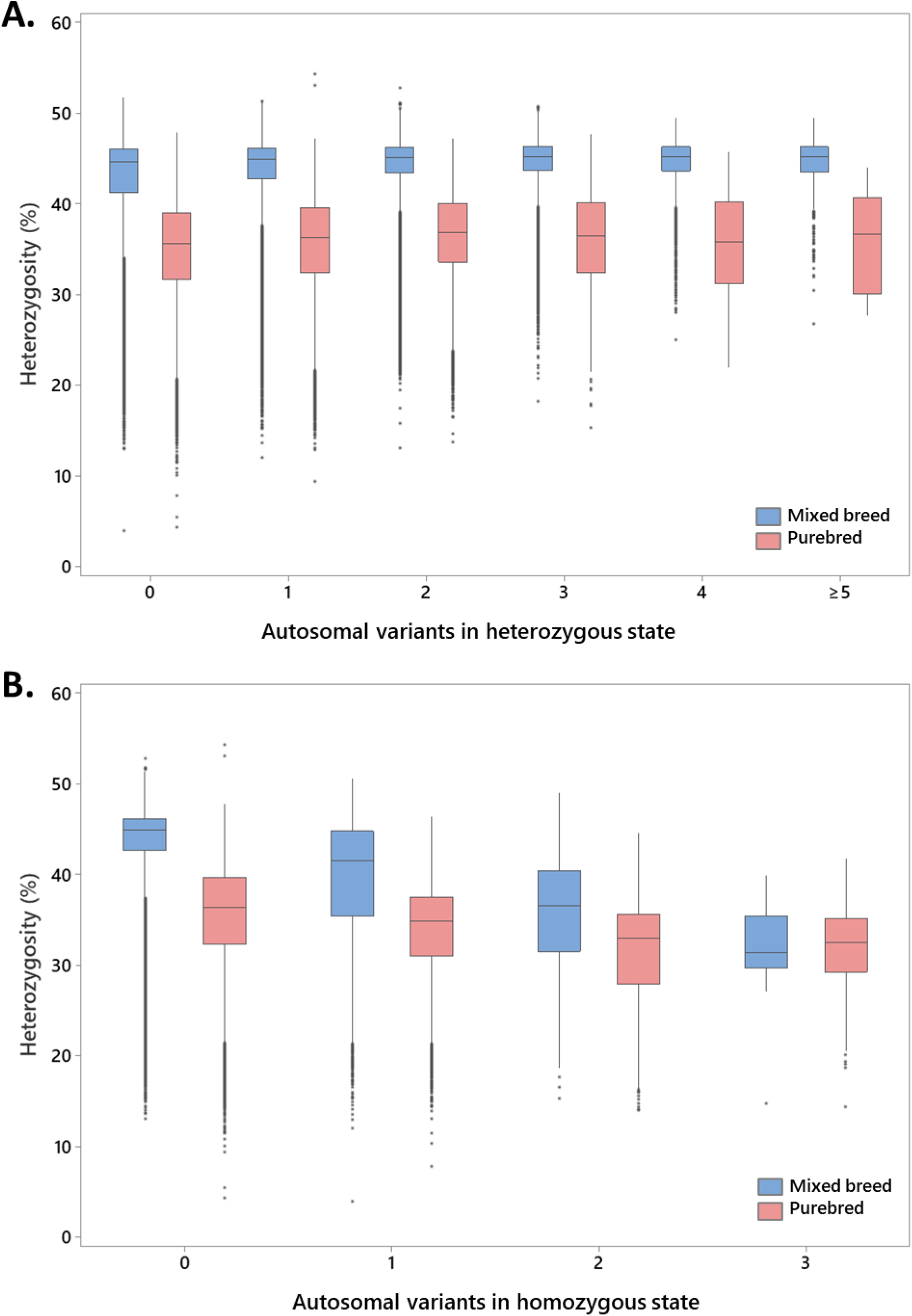
The relationship between genome-wide genetic heterozygosity and autosomal disease variants present in A) heterozygous state and B) homozygous state; based on 178 recessive and 13 semi-dominant or dominant variants tested in 1,054,293 dogs. Medians and interquartile range boxes representing the middle 50% and whiskers representing the bottom and top 25% of the data excluding outliers (data points >1.5x the interquartile range from the boxes; asterisks) are shown.

### The most common disease variants in dogs

The top 20% of the most prevalent disease alleles collectively accounted for 98.3% of all disease-associated alleles observed in the study sample, and the distribution was highly similar across both the mixed breed dogs and combined purebred sample groups (Table 1; full details provided as Supplementary Table 1). Only 7 of the 50 most common variants in mixed breed dogs were not present among the purebred top 50 variants, and vice versa. Among the most common variants were several variants with a suggested ancient origin and established distribution across a wide range of modern-day breeds, such as chondrodystrophy and intervertebral disc disease (CDDY), degenerative myelopathy (DM), progressive rod-cone degeneration (prcd-PRA), hyperuricosuria (HUU) and collie eye anomaly (CEA) (12–16). On the other hand, we observed a high frequency of several variants that have so far been linked to specific disease phenotypes based on examinations of a single breed or subpopulation within a breed including *IGFBP5* gene candidate variant for bald thigh syndrome in sighthounds (17), *SLC7A9* and *SLC3A1* gene variants associated with cystinuria in Bulldogs (18, 19), as well as *PDK4* and *TTN* gene risk variants for dilated cardiomyopathy (DCM) in Dobermans (20–22). The clinical impact of these broadly distributed putative genetic risk factors in dogs of a diverse genetic ancestry background were further assessed with the use of electronic medical records.

**Table 1.** Top 50 most frequently observed genetic variants in the examined canine population.

| OMIA variant ID<sup>1</sup> | Variant phenotype | All dogs |  | Mixed Breed Dogs |  | Purebred Dogs |  |
| --- | --- | --- | --- | --- | --- | --- | --- |
|  |  | Prevalence Rank | Allele frequency [%] | Prevalence Rank | Allele frequency [%] | Prevalence Rank | Allele frequency [%] |
| 855 | Chondrodystrophy and Intervertebral Disc Disease Risk (CDDY) | 1 | 11.951 | 1 | 11.353 | 1 | 13.945 |
| 36 | Degenerative Myelopathy (DM) | 2 | 7.858 | 2 | 7.506 | 2 | 9.037 |
| 699 | Cone-Rod Dystrophy (cord1-PRA/crd4) | 3 | 3.809 | 3 | 3.641 | 3 | 4.370 |
| 76 | Progressive Rod-Cone Degeneration (prcd-PRA) | 4 | 2.923 | 4 | 3.314 | 8 | 1.613 |
| 002168-9615 | Bald Thigh Syndrome (Discovered in Sighthounds) | 5 | 2.373 | 5 | 2.742 | 10 | 1.358 |
| 000256-9615 | Cystinuria Type I-B (SLC7A9 p.A217T) | 6 | 1.991 | 7 | 1.603 | 4 | 3.291 |
| 83 | Hyperuricosuria (HUU) | 7 | 1.873 | 6 | 1.841 | 7 | 1.982 |
| 000256-9615 | Cystinuria Type I-A (SLC3A1 p.I192V) | 8 | 1.739 | 8 | 1.372 | 5 | 2.969 |
| 000256-9615 | Cystinuria Type I-A (SLC3A1 p.S698G) | 9 | 1.516 | 11 | 1.118 | 6 | 2.847 |
| 000162-9615 | Dilated Cardiomyopathy risk factor (Discovered in the Doberman Pinscher; PDK4-related) | 10 | 1.272 | 9 | 1.258 | 11 | 1.321 |
| 632 | Collie Eye Anomaly (CEA) | 11 | 1.083 | 10 | 1.141 | 15 | 0.888 |
| 39 | Exercise-Induced Collapse (EIC) | 12 | 0.951 | 14 | 0.923 | 12 | 1.046 |
| 000162-9615 | Dilated Cardiomyopathy risk factor (Discovered in the Doberman Pinscher; TTN-related) | 13 | 0.947 | 12 | 1.022 | 18 | 0.698 |
| 469 | MDR1 (Multidrug Resistance 1) Medication Sensitivity | 14 | 0.926 | 13 | 0.974 | 17 | 0.763 |
| 616 | Ichthyosis (Discovered in the Golden Retriever) | 15 | 0.910 | 16 | 0.734 | 9 | 1.500 |
| 401 | Von Willebrand's Disease, Type 1 (vWD 1) | 16 | 0.776 | 15 | 0.736 | 14 | 0.913 |
| 275 | Canine Multifocal Retinopathy 1 (Discovered in Mastiff-related breeds; CMR1) | 17 | 0.701 | 17 | 0.607 | 13 | 1.013 |
| 1050 | Stargardt Disease (Discovered in the Labrador Retriever) | 18 | 0.626 | 18 | 0.576 | 16 | 0.791 |
| 365 | Primary Lens Luxation (PLL) | 19 | 0.455 | 19 | 0.563 | 34 | 0.094 |
| 67 | Neuronal Ceroid Lipofuscinosis 4A (Discovered in the American Staffordshire Terrier; NCL4A) | 20 | 0.451 | 21 | 0.405 | 19 | 0.607 |
| 000247-9615 | Pituitary-Dependent Hyperadrenocorticism (Discovered in Poodles) | 21 | 0.364 | 20 | 0.418 | 25 | 0.186 |
| 40 | Factor VII Deficiency | 22 | 0.356 | 22 | 0.361 | 20 | 0.341 |
| 74 | Prekallikrein Deficiency | 23 | 0.317 | 23 | 0.310 | 21 | 0.340 |
| 002203-9615 | Ehlers-Danlos Syndrome (Discovered in the Chihuahua and Poodle) | 24 | 0.283 | 25 | 0.295 | 22 | 0.252 |
| 1083 | Progressive Retinal Atrophy (Discovered in the Giant Schnauzer; *NECAP1*-related) | 25 | 0.255 | 24 | 0.306 | 37 | 0.083 |
| 112 | Shar-Pei Autoinflammatory Disease (SPAID) | 26 | 0.216 | 26 | 0.244 | 30 | 0.123 |
| 49 | Hypocatalasia | 27 | 0.196 | 27 | 0.197 | 24 | 0.194 |
| 528 | Cone-Rod Dystrophy 1 (Discovered in the American Staffordshire Terrier; crd1) | 28 | 0.164 | 28 | 0.161 | 27 | 0.177 |
| 949 | Progressive Retinal Atrophy (Discovered in the Golden Retriever; GR_PRA2) | 29 | 0.145 | 30 | 0.124 | 23 | 0.217 |
| 422 | Canine Scott Syndrome (CSS) | 30 | 0.126 | 29 | 0.135 | 33 | 0.095 |
| 280 | Persistent Müllerian Duct Syndrome (PMDS) | 31 | 0.112 | 34 | 0.091 | 26 | 0.182 |
| 78 | Skeletal Dysplasia 2 (SD2) | 32 | 0.101 | 31 | 0.104 | 36 | 0.088 |
| 411 | Craniomandibular Osteopathy (Discovered in the Cairn, Scottish and West Highland White Terrier) | 33 | 0.096 | 35 | 0.087 | 29 | 0.127 |
| 563 | Ichthyosis (Discovered in the American Bulldog; *NIPAL4*-related) | 34 | 0.087 | 32 | 0.096 | 43 | 0.058 |
| 924 | Centronuclear Myopathy (Discovered in the Labrador Retriever; *HACD1*-related; CNM) | 35 | 0.080 | 36 | 0.081 | 39 | 0.074 |
| 001326-9615 | Protein Losing Nephropathy (PLN; *NPHS1*-related) | 36 | 0.077 | 40 | 0.064 | 31 | 0.123 |
| 86 | Hereditary Nasal Parakeratosis (Discovered in the Labrador Retriever; HNPK) | 37 | 0.076 | 43 | 0.057 | 28 | 0.140 |
| 527 | Cystinuria Type II-A (Discovered in the Australian Cattle Dog) | 38 | 0.075 | 33 | 0.093 | 74 | 0.013 |
| 995 | Neuroaxonal Dystrophy (Discovered in the Rottweiler; *VPS11*-related) | 39 | 0.073 | 38 | 0.077 | 42 | 0.062 |
| 447 | Intestinal Cobalamin Malabsorption (Discovered in the Border Collie; *CUBN*-related) | 40 | 0.071 | 42 | 0.061 | 32 | 0.102 |
| 804 | Congenital Myasthenic Syndrome (Discovered in the Heideterrrier; *CHRNE*-related; CMS) | 41 | 0.068 | 37 | 0.079 | 48 | 0.036 |
| 642 | Osteochondrodysplasia (Discovered in the Miniature Poodle) | 42 | 0.055 | 39 | 0.065 | 56 | 0.023 |
| 546 | Polyneuropathy with Ocular Abnormalities and Neuronal Vacuolation (Discovered in the Black Russian Terrier and Rottweiler; POANV) | 43 | 0.054 | 48 | 0.051 | 40 | 0.065 |
| 942 | Primary Open Angle Glaucoma and Lens Luxation (Discovered in the Chinese Shar-Pei) | 44 | 0.054 | 41 | 0.062 | 51 | 0.028 |
| 606 | Cone-Rod Dystrophy 2 (Discovered in the Pit Bull Terrier; crd2) | 45 | 0.053 | 47 | 0.052 | 44 | 0.056 |
| 98 | Macrothrombocytopenia (Discovered in the Norfolk and Cairn Terrier) | 46 | 0.049 | 45 | 0.055 | 53 | 0.027 |
| 1323 | Dilated Cardiomyopathy (Discovered in the Schnauzer; DCM) | 47 | 0.048 | 44 | 0.056 | 58 | 0.022 |
| 478 | Trapped Neutrophil Syndrome (TNS) | 48 | 0.048 | 52 | 0.039 | 38 | 0.078 |
| 575 | Progressive Retinal Atrophy (Discovered in the Golden Retriever; GR_PRA1) | 49 | 0.048 | 53 | 0.036 | 35 | 0.088 |
| 444 | Acral Mutilation Syndrome (AMS) | 50 | 0.047 | 50 | 0.050 | 46 | 0.037 |

<sup>1</sup> OMIA (Online Mendelian Inheritance in Animals; omia.org) ID number where a specific variant ID was not available at the time of writing

### Uncovering the breed distribution of canine genetic variants

Among the main factors shaping the disease heritage of the purebred dog population are breed formation, or founding, events that are typically coupled with closing of the breed registry to limit additional gene flow to the novel breed. Later crossbreeding can transfer disease variants from one breed to another as we have shown earlier (6), while mutations arising within breeds further shape the gene pool. While the presence or absence of a disease variant in a specific breed background can be understood in the context of the aforementioned breeding practices, their full distribution across breeds remains elusive without extensive genetic screening in large and diverse sample sets. The detailed breed distributions and allele frequencies of all variants tested in the present study, including the aforementioned high frequency variants in the *IGFBP5*, *SLC7A9*, *SLC3A1*, *PDK4* and *TTN* genes, are summarized and available as a resource for the community in Supplementary Table 3. For the purposes of this study, we defined a novel breed finding of particular interest as a disease variant present in an additional pure breed with an allele frequency of ≥1%, with the additional requirement of more than one carrier/affected dog observed. With this definition, we discovered the genetic presence of 26 disease variants in a total of 65 pure breeds in which they, to the best of our knowledge, have not been characterized previously in the peer-reviewed scientific literature (Table 2).

**Table 2.** Disease variants found in additional breeds at an allele frequency ≥1%

| OMIA Variant ID<sup>1</sup> | Variant phenotype | Previously identified breed(s) | Additional identified breed(s) | Total N dogs screened | Variant frequency (%) |
| --- | --- | --- | --- | --- | --- |
| 444 | Acral Mutilation Syndrome (AMS) | English Pointer, English Springer Spaniel, French Spaniel, German Shorthaired Pointer | Cirneco dell'Etna | 7 | 14,29 |
|  |  |  | Irish Terrier | 35 | 2,86 |
|  |  |  | Old English Sheepdog | 423 | 7,92 |
| 1044 | Amelogenesis Imperfecta (Discovered in the Parson Russell Terrier; AI) | Parson Russell Terrier | Danish Swedish Farmdog | 61 | 3,28 |
| 275 | Canine Multifocal Retinopathy 1 (Discovered in Mastiff-related breeds; CMR1) | Boerboel, Bull Mastiff, English Mastiff, Great Pyrenees | Boston Terrier | 3702 | 7,47 |
|  |  |  | Neapolitan Mastiff | 90 | 1,11 |
| 422 | Canine Scott Syndrome (CSS) | German Shepherd Dog | Presa Canario | 64 | 1,56 |
| 632 | Collie Eye Anomaly (CEA) |  | Anatolian Shepherd Dog | 66 | 3,79 |
|  |  |  | Lacy Dog | 32 | 9,38 |
|  |  |  | Maremma Sheepdog | 37 | 2,7 |
|  |  |  | McNab | 28 | 17,86 |
| 699 | Cone-Rod Dystrophy (cord1-PRA/crd4) | Miniature Long-haired Dachshund | Airedale Terrier | 200 | 3,5 |
|  |  |  | American Eskimo Dog | 302 | 3,48 |
|  |  |  | American Staffordshire Terrier | 42793 | 17,75 |
|  |  |  | Australian Shepherd | 2296 | 1,57 |
|  |  |  | Barbet | 106 | 5,66 |
|  |  |  | Basset Hound | 990 | 1,32 |
|  |  |  | Beagle | 5292 | 10,51 |
|  |  |  | Bedlington Terrier | 11 | 9,09 |
|  |  |  | Bloodhound | 280 | 21,79 |
|  |  |  | Catahoula Leopard Dog | 154 | 4,25 |
|  |  |  | Caucasian Shepherd Dog | 41 | 2,44 |
|  |  |  | Chihuahua | 4273 | 3,88 |
|  |  |  | Clumber Spaniel | 12 | 37,5 |
|  |  |  | English Cocker Spaniel | 580 | 1,73 |
|  |  |  | Field Spaniel | 29 | 12,07 |
|  |  |  | French Bulldog | 13114 | 7,96 |
|  |  |  | Kai Ken | 9 | 33,33 |
|  |  |  | Lagotto Romagnolo | 623 | 4,27 |
|  |  |  | Poodle (Miniature) | 3555 | 1,13 |
|  |  |  | Pumi | 86 | 4,65 |
|  |  |  | Rottweiler | 4718 | 6,82 |
|  |  |  | Schnauzer (Standard) | 75 | 12,67 |
|  |  |  | Silky Terrier | 28 | 8,93 |
|  |  |  | Weimaraner | 647 | 6,66 |
| 1037 | Congenital<br>Dyshormonogenic<br>Hypothyroidism with<br>Goiter (Discovered in the<br>Shih Tzu) | Shih-Tzu | Pekingese | 239 | 1,05 |
| 1079 | Deafness and Vestibular<br>Dysfunction (Discovered<br>in the Doberman<br>Pinscher) | Doberman Pinscher | Mudi | 42 | 4,76 |
| 002203-9615 | Ehlers-Danlos Syndrome<br>(Discovered in the<br>Chihuahua and Poodle) | Chihuahua, Poodle | German Pinscher | 13 | 38,46 |
|  |  |  | German Shorthaired<br>Pointer | 65 | 2,31 |
|  |  |  | Pomeranian | 121 | 4,55 |
| 39 | Exercise-Induced Collapse (EIC) | Chesapeake Bay Retriever, Curly-coated retriever, Labrador Retriever | Maltese | 2413 | 2,15 |
| 40 | Factor VII Deficiency | Beagle | Italian Greyhound | 263 | 1,33 |
|  |  |  | Lacy Dog | 32 | 10,94 |
|  |  |  | Schipperke | 72 | 9,03 |
| 970 | Hereditary Nasal Parakeratosis (Discovered in the Greyhound; HNPK) | Greyhound | Saluki | 24 | 4,17 |
| 83 | Hyperuricosuria (HUU) | Dalmatian | Argentine Dogo | 225 | 2 |
|  |  |  | Catahoula Leopard Dog | 154 | 1,95 |
|  |  |  | German Wirehaired Pointer | 84 | 1,79 |
|  |  |  | Munsterlander (Small) | 15 | 10 |
| 49 | Hypocatalasia | American Foxhound, Beagle | Plott | 25 | 8 |
| 98 | Macrothrombocytopenia (Discovered in the Norfolk and Cairn Terrier) | Norfolk Terrier | Caucasian Shepherd Dog | 41 | 2,44 |
|  |  |  | Chesapeake Bay Retriever | 138 | 3,99 |
| 469 | MDR1 (Multidrug Resistance 1) Medication Sensitivity | Australian Shepherd, Border Collie, Collie, German Shepherd Dog, Longhaired Whippet, Miniature Australian Shepherd, Old English Sheepdog, Shetland Sheepdog, Silken Windhound, Waller, White Swiss Shepherd | Lacy Dog | 32 | 1,56 |
| 993 | Microphthalmia (Discovered in the Soft-Coated Wheaten Terrier) | Irish Soft-Coated Wheaten Terrier | Russell Terrier | 239 | 6,49 |
| 69 | Neuronal Ceroid Lipofuscinosis 8 (Discovered in the English Setter; NCL8) | English Setter | Gordon Setter | 20 | 5 |
| 642 | Osteochondrodysplasia (Discovered in the Miniature Poodle) | Miniature Poodle | Bichon Frise | 1069 | 2,43 |
| 000247-9615 | Pituitary-Dependent Hyperadrenocorticism (Discovered in Poodles) | Poodle | Australian Cattle Dog | 982 | 2,6 |
|  |  |  | Great Pyrenees | 1985 | 11,79 |
|  |  |  | Maremma Sheepdog | 37 | 5,41 |
|  |  |  | Miniature American Shepherd | 1476 | 2,24 |
|  |  |  | Portuguese Podengo Pequenos | 17 | 11,76 |
|  |  |  | Spanish Water Dog | 96 | 8,85 |
| 74 | Prekallikrein Deficiency | Shih-Tzu | Lhasa Apso | 243 | 3,5 |
| 365 | Primary Lens Luxation (PLL) | Jack Russell Terrier, Lancashire Heeler, Miniature Bull Terrier | Coyote | 9 | 16,67 |
|  |  |  | McNab | 28 | 7,14 |
|  |  |  | Scottish Terrier | 237 | 1,05 |
| 1391 | Bardet-Biedl syndrome 2 or Progressive Retinal Atrophy (Discovered in the Shetland Sheepdog; BBS2-PRA) | Shetland Sheepdog | Miniature American Shepherd | 1476 | 1,36 |
| 76 | Progressive Rod-Cone Degeneration (prcd-PRA) | American Cocker Spaniel, Australian Cattle Dog, Australian Shepherd, Australian Stumpy Tail Cattle Dog, Chesapeake Bay Retriever, Chinese Crested Dog, English Cocker Spaniel, Entlebucher Mountain Dog, Finnish Lapphund, Golden Retriever, Karelian Bear Dog, Kuvasz, Labrador Retriever, Lapponian Herder, Miniature Poodle, Norwegian Elkhound, Nova Scotia Duck Tolling Retriever, Portuguese Water Dog, Spanish Water Dog, Swedish Lapphund, Toy Poodle, Yorkshire Terrier | Bichon Frise | 1069 | 1,13 |
|  |  |  | Dalmatian | 820 | 2,39 |
|  |  |  | Kai Ken | 9 | 11,11 |
|  |  |  | Schnauzer (Miniature) | 4638 | 1,11 |
| 552 | Van den Ende-Gupta Syndrome (VDEGS) | Wirehaired Fox Terrier | Welsh Terrier | 21 | 4,76 |
| 401 | Von Willebrand's Disease, Type 1 (vWD 1) | Doberman Pinscher, Kromfohrlander | Boerboel | 165 | 1,21 |
|  |  |  | Keeshond | 70 | 4,29 |
|  |  |  | Lacy Dog | 32 | 1,56 |
|  |  |  | Maltese | 2413 | 2,03 |

<sup>1</sup> OMIA (Online Mendelian Inheritance in Animals; omia.org) ID number where a specific variant ID was not
available at the time of writing

### Mining of electronic medical records (EMRs) establishes clinical significance of disease variants and uncovers evidence for screening relevance on additional breed ancestry backgrounds

*A*dditional evidence supporting variant clinical significance was collected to strengthen the current body of evidence supporting pathogenicity of selected variants, with a special focus on establishing the relevance of unexpected variant findings (variants not previously reported to be present in the breed based on the scientific literature). Medical records were available for a significant proportion of dogs of the genotyped cohort (43.5%; N = 458,433) that had attended a Banfield Pet Hospital^Ⓡ^ veterinary clinic for consultation or treatment, and additional direct interviews with owners were carried out to obtain supplemental phenotype information where possible. With this approach, we evaluated the phenotypes of 12028 dogs that were genetically at risk based on their DNA testing result for one of 49 inherited disease variants (Table 3; full details by dog in Supplementary Table 4). In particular, among the variants evaluated in more than one genetically affected dog, we observed complete penetrance (defined as 100% of dogs genetically at risk showing clinical signs of the expected disease) of the following variants: canine leukocyte adhesion deficiency (CLAD) Type III, disproportionate short-limbed chondrodysplasia (discovered in the Norwegian Elkhound; *ITGA10*-related), cone-rod dystrophy 2 (discovered in the Pit Bull Terrier; crd2), focal non-epidermolytic palmoplantar keratoderma (discovered in the Dogue de Bordeaux), hemophilia A (discovered in German Shepherd Dog; *F8* p.C548Y variant), hereditary footpad hyperkeratosis (discovered in the Irish Terrier and Kromfohrländer), ichthyosis (discovered in the American Bulldog; *NIPAL4*-related), neuroaxonal dystrophy (discovered in the Rottweiler; *VPS11*-related), skeletal dysplasia 2 (SD2), and trapped neutrophil syndrome (TNS).

**Table 3.**
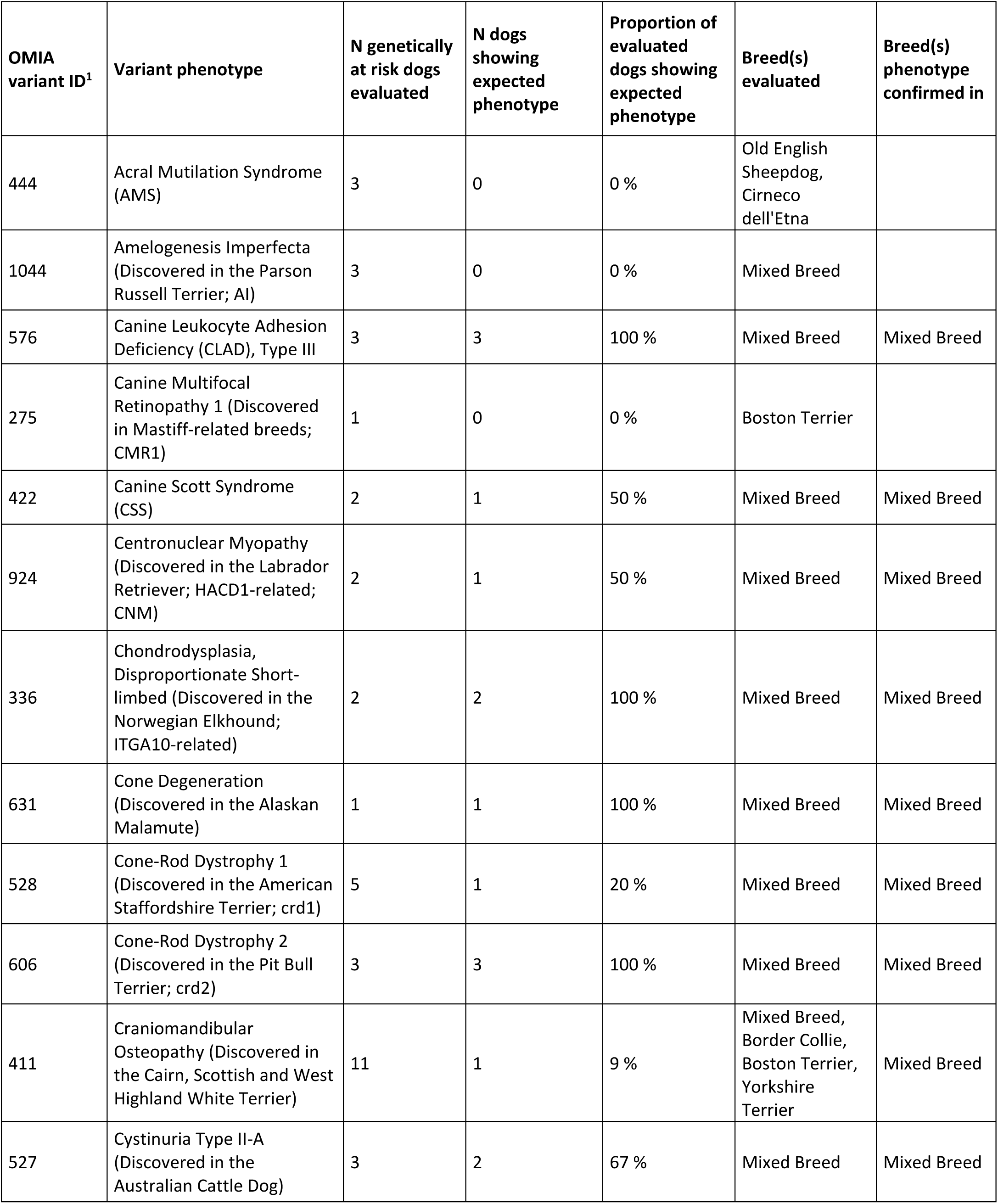
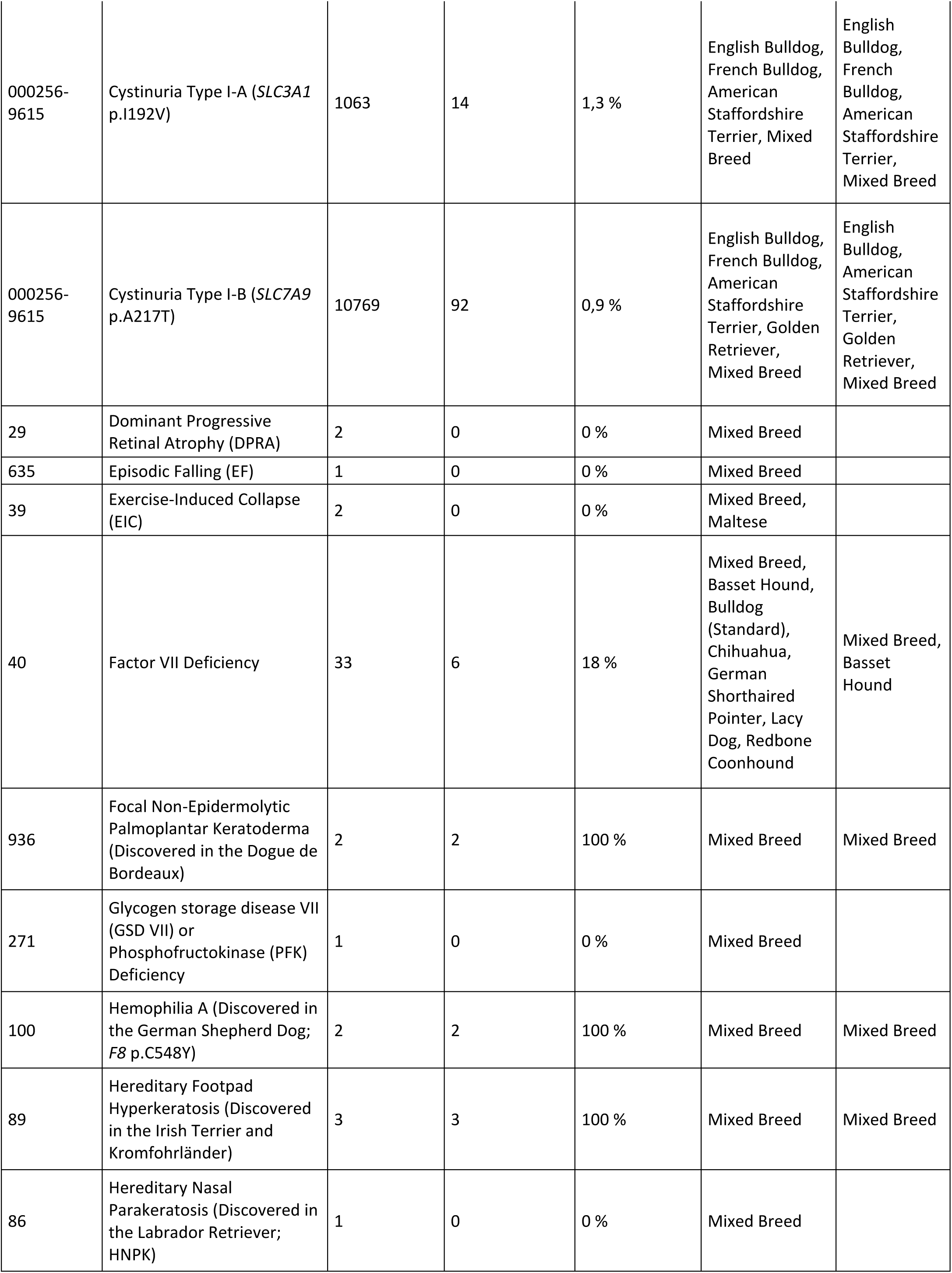
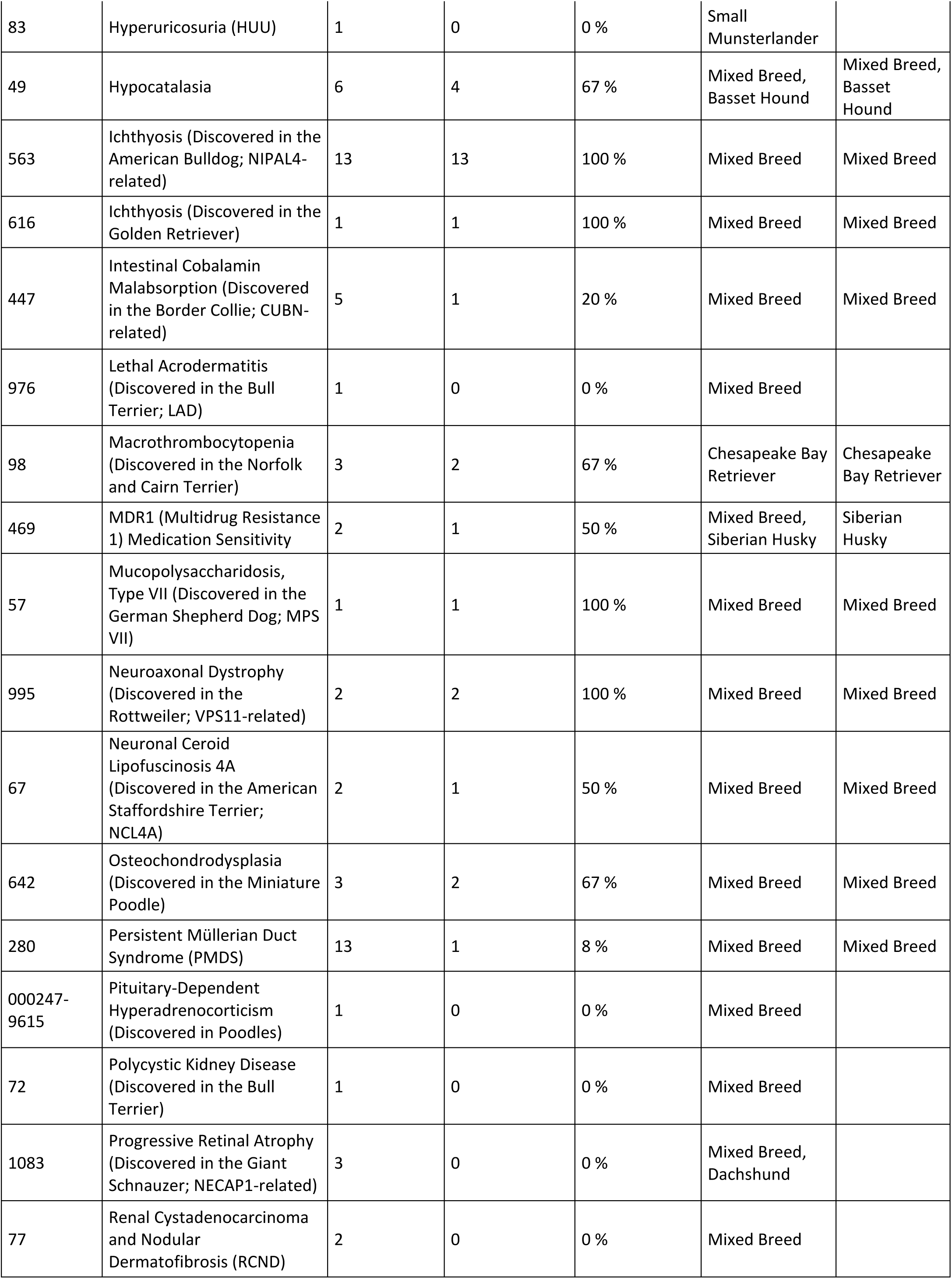
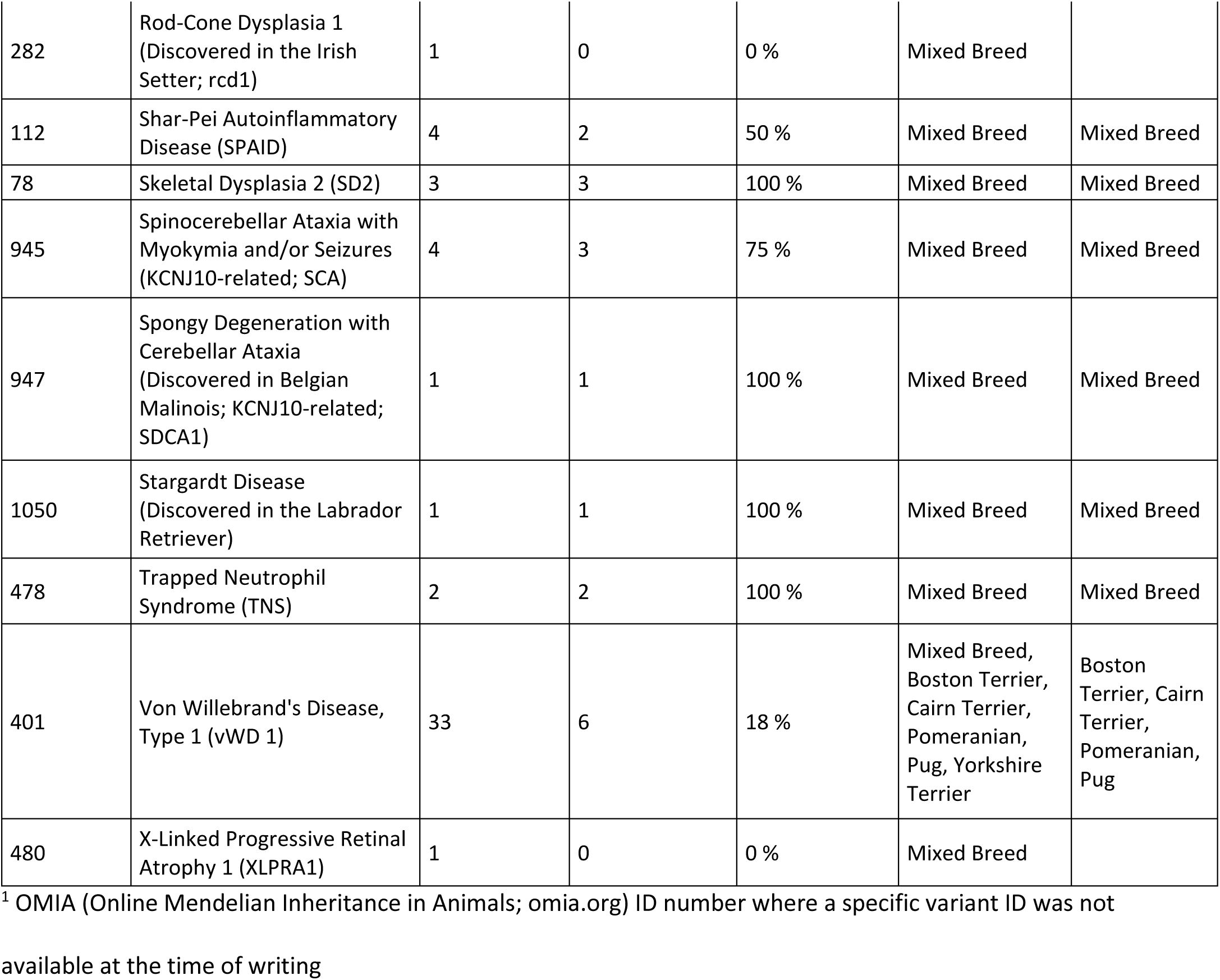
Summary of clinical validation examinations

Overall, the additional compiled evidence supporting causality of the variants listed above was obtained on diverse genetic backgrounds, with varying identified contributions of ancestry from the original variant discovery breeds (Supplementary Table 4). With focus on variants findings not previously documented in the scientific literature in specific purebreds, we found plausible evidence (based on genetically at risk dogs showing at least some clinical signs that are known manifestations of the evaluated disease) for the clinical relevance of cystinuria in the French Bulldog, factor VII deficiency in the Basset Hound, hypocatalasia in the Basset Hound, macrothrombocytopenia in the Chesapeake Bay Retriever, multidrug resistance 1 (MDR1) related medication sensitivity in the Siberian Husky, and von Willebrand’s Disease Type 1 (vWD 1) in the Boston Terrier, Cairn Terrier, Pomeranian and Pug. Selected validation observations are presented in more detail in the following sections.

### Inherited catalase deficiency is a notable cause of oral ulcers and periodontal disease

The clinical ramifications of the *CAT* gene variant associated with hypocatalasia (23) were not well characterized in the pet population previously, as the disorder has been studied mainly in a laboratory colony of Beagles. Using EMR data, we were able to confirm signs consistent with hypocatalasia with an early age of onset in mixed breed dogs with hound ancestry, as well as in one purebred Basset Hound and a mixed breed dog without recent scent hound ancestry (Table 3; Supplementary Table 4). Clinical signs of the disorder were fairly consistent across dogs at genetic risk, and generally included areas of oral ulceration along with gingival recession with bone loss/necrosis and tooth root exposure, causing pain, loss of teeth, and in some cases prolonged bleeding from the mouth. Oral repairs had a tendency to fail due to friable gingival tissue, and clinical signs were progressive in all pets reviewed. These observations, together with our reports of genetic presence of the hypocatalasia variant in pet Beagles, American and English Foxhounds, Harrier, Treeing Walker Coonhound, and Plott Hounds ((6, 7) and the present study) call for greater awareness of hypocatalasia as a health concern especially in dogs of scent hound ancestry.

### Factor VII deficiency presents as a common subclinical propensity for prolonged blood coagulation

The number of identified breeds harboring the *F7* gene variant associated with Factor VII deficiency (24) has significantly increased after the introduction of genetic panel screening technologies, with the current total exceeding 25 breeds and breed varieties of diverse ancestry (4,6,7). In the present study, EMRs included clotting test laboratory result values for four dogs (three mixed breed dogs and one Basset Hound) homozygous for the *F7* variant. All had an elevated prothrombin time (PT) and a normal activated partial thromboplastin time (aPTT) as anticipated (Supplementary Table 4). One of the mixed breed dogs had additionally received factor VII testing, and the value was reduced as expected at 35% of baseline levels. Two additional mixed breed dogs homozygous for the *F7* variant did not have recorded blood clotting measurements but had sought veterinary care due to episodes of hematochezia and hematemesis. Taken together, we note that all blood clotting measurements from *F7* homozygous dogs available for examination by the authors to date have shown the same pattern of elevated PT but normal aPTT times ((6) and the present study). These observations further strengthen the perception that Factor VII deficiency manifests as a mild propensity for prolonged blood clotting, that typically remains clinically undetected, in any dog genetically at risk, regardless of genetic ancestry.

### Multiple genetic causes of a “short-legged” phenotype in dogs

Related retrogenes on chromosomes 12 (12-FGF4RG) and 18 (18-FGF4RG) represent common and widespread genetic causes of disproportionate dwarfism in dogs, in the form of a characteristic short-limbed phenotype that has become a hallmark of the breed standard in breeds such as Dachshunds, Corgis, and Basset Hounds (12, 25). The chromosome 12 retrogene 12-FGF4RG has particular health relevance as it simultaneously confers increased risk for intervertebral disc disease (IVDD) and is associated with age at time of surgical treatment across mixed breed dogs and all affected breeds (26). We confirm the ubiquitous nature of the 12-FGF4RG variant as the most common tested variant observed in dogs in the present study, with an allele frequency of 11.4% in mixed breed dogs and 13.9% in the combined purebred dog sample (Table 1) and presence in 89 of the examined breeds or breed varieties at a frequency ≥1% (Supplementary Data).

In addition to the aforementioned widespread variants for short-leggedness that are inherited in an autosomal dominant or semi-dominant manner, other rare recessive variants have been identified as well. We provide additional confirmation of the causal effect of two such variants, skeletal dysplasia 2 (SD2; also known as mild disproportionate dwarfism) and *ITGA10* gene-related disproportionate short-limbed chondrodysplasia (27, 28), on mixed breed ancestry backgrounds. The three dogs homozygous for SD2 were all genotyped as being devoid of any copies of either 12-FGF4RG or 18-FGF4RG, and described in their medical history as displaying “all short limbs”, “shortened legs and body longer than dog is tall”, being “cow-hocked” or having a carpal valgus deformity (Figure 4A). Notably, none of the phenotypically evaluated dogs tested as having more than around 5% ancestry contribution from the original variant discovery breed, the Labrador Retriever. The two mixed breed dogs homozygous for the *ITGA10* were described as having shortened legs with a normal sized head and body, slight bowing or crookedness of the front legs, and one of them in addition displayed the characteristic abnormally short outer digits previously associated with this phenotype (Figure 4B-C). Neither of the two showed a notable genetic ancestry contribution (≥1%) from the breeds in which the effects of the *ITGA10* variant have been characterized to date (the Norwegian Elkhound, the Karelian Bear Dog, and the Chinook (6, 27)).

**Figure 4.**
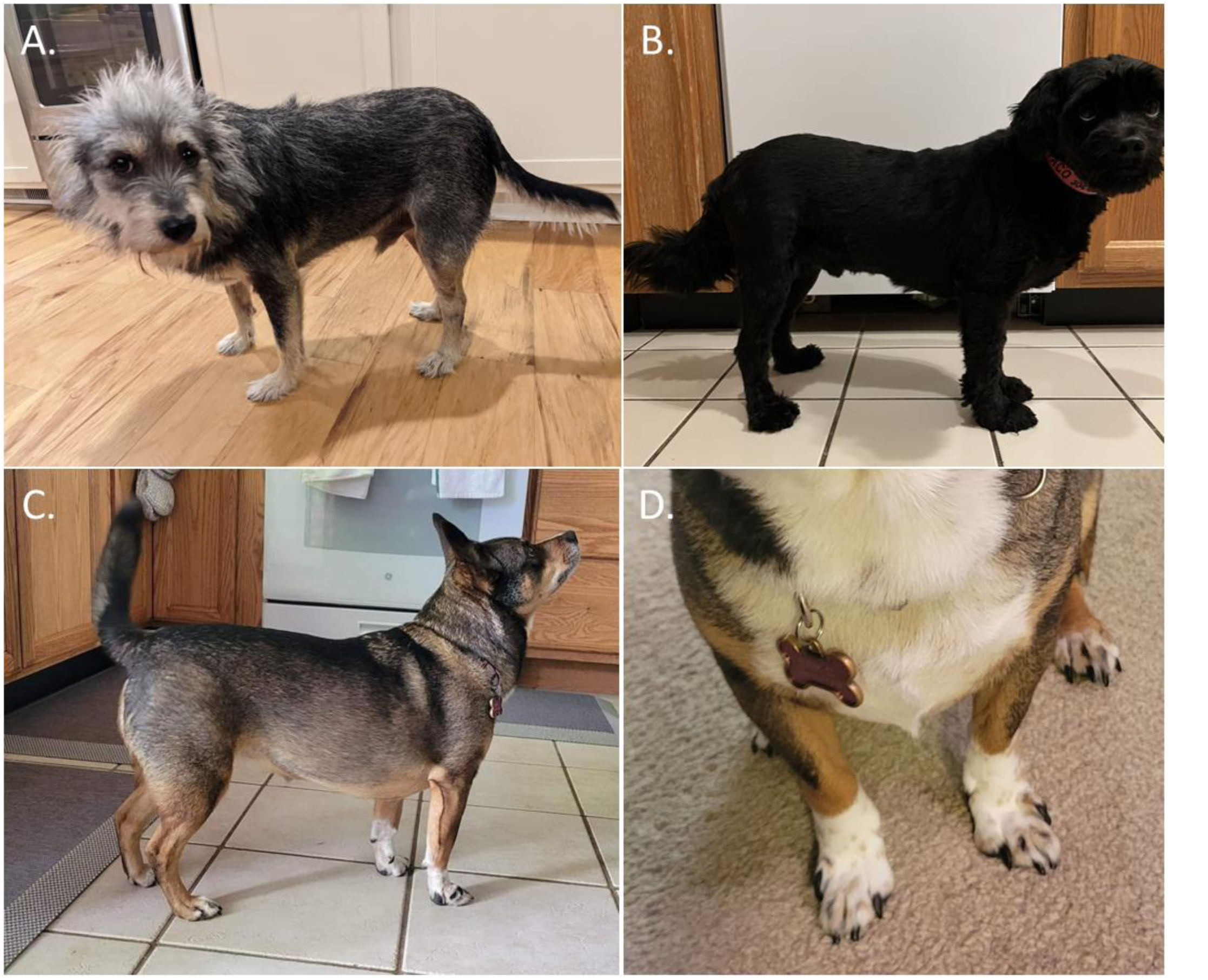
Disproportionate dwarfism phenotypes in mixed breed dogs. A) Skeletal dysplasia 2 (SD2) manifests as a mild phenotype in which the front legs are typically shorter than the hind legs and the body and head are of normal size. B-C) *ITGA10* gene-related disproportionate short-limbed chondrodysplasia manifests as shortened, typically crooked legs with a normal sized head and body, and in some cases like D) also as abnormally short outer digits.

### Genetic and phenotypic heterogeneity underlying canine cystinuria

To date, studies on canine cystinuria have implicated involvement of variants in the functional candidate genes *SLC3A1* and *SLC7A9*, encoding amino acid transporters involved in the excretion of cystine and known to explain approximately 70% of human cystinuria cases (18, 29). Among the putative variants contributing to cystinuria we selected for genotyping in the present study were two missense variants (*SLC3A1* p.I192V and p.S698G) previously implicated in a recessive form of cystinuria affecting English Bulldogs and potentially also French Bulldogs, and a missense variant *SLC7A9* p.A217T discovered present in the heterozygous state in one English Bulldog diagnosed with cystinuria (18, 19). We find that the aforementioned variants are present in more than 30 breeds of diverse ancestry with some of the highest allele frequencies observed in Bulldog and Mastiff-type breeds (Supplementary Table 4). While the SLC3A1 variants coexist in full linkage disequilibrium in many, but not all, breeds they are present in, we observed several non-mastiff breeds with a high frequency of the SLC7A9 variant alone (e.g., Golden Retriever, Swedish Vallhund, and English Cocker Spaniel; allele frequencies 12-26%). Within the scope of this study, we focused on clinically validating the effect of the SLC3A1 p.I192V and p.S698G gene variants in mixed breed dogs and in the three breeds in which the largest number of more than 1-year-old alternative allele homozygotes were found and the two variants were genetically linked: the English/Standard Bulldog, the French Bulldog, and the American Staffordshire Terrier. Only 6/400 (1.5%) English Bulldogs, 5/424 French Bulldogs (1.2%), 0/39 American Staffordshire Terriers (0%), and 3/200 (1.5%) mixed breed dogs at putative genetic risk showed clinical signs (EMR-documented cystic calculi, crystalluria, hematuria, urolithiasis, nephrolithiasis, urinary tract infection or cystitis) supporting a diagnosis of cystinuria. The outcome was similar when dogs from the same breeds, plus the Golden Retriever (examined due to a high 24% breed allele frequency), were evaluated for the SLC7A9 p.A217T variant. Only 17/1872 (0.9%) English Bulldogs, 0/69 (0%) French Bulldogs, 5/691 (0.7%) American Staffordshire Terriers, 26/2571 (1.0%) Golden Retrievers and 44/5522 (0.8%) mixed breed dogs at putative genetic risk due to presence of either one or two copies of SLC7A9 p.A217T had EMRs with plausible evidence of cystinuria. As these values are likely underestimates of disease risk due to undiagnosed signs of cystinuria, gender-specific differences in manifestation, or later onset of disease, we cannot exclude the possibility that the studied missense variants represent modifiers or linked markers predictive of cystinuria risk on some breed backgrounds such as Bulldog ancestry. Given the widespread nature of the examined variants across multiple breeds it is however likely that they are non-causal in themselves, and further studies are warranted to identify additional risk factors for cystinuria and ensure appropriate genetic counseling.

In contrast to the aforementioned form of cystinuria, another variant in the *SLC3A1* gene (p.T366_T367del) was found to cause an autosomal dominant form of the disease in Australian Cattle Dogs, with a more severe phenotype in homozygous dogs (29). We evaluated three mixed breed dogs, two homozygous and one heterozygous for *SLC3A1* p.T366_T367del, and confirm that the variant was associated with a fairly severe morbidity with the examined homozygous patients showing their first signs of obstruction by 7 months of age. Both affected dogs passed or were euthanized before the age of two years. The third dog heterozygous for the variant and with available EMR data has not returned to the clinic since approximately nine months of age but did not show any significant clinical diagnoses except for submissive urination at the last visit.

### Additional evidence for specific highly penetrant and clinically relevant canine disease variants

In addition, the EMRs provided further evidence supporting the causality and high penetrance of variants previously associated with cone-rod dystrophy 2 (crd2) (30), spinocerebellar ataxia with myokymia and/or seizures (SCA; *KCNJ10*-related) (31), and both the Golden Retriever (*PNPLA1*-related) (32) and American Bulldog (*NIPAL4*-related) ichthyoses (33). In the three mixed breed dogs evaluated due to their expected genetic predisposition to develop crd2, the retinal condition appears to show a particularly early age of onset, with overt clinical signs beginning at 3-5 months of age. Evaluated dogs were all reported to show signs of vision loss such as walking into objects, loss of the menace response, and clumsiness, and all were apparently blind on exam by the age of one year. Similarly, the genetic variant associated with SCA (31) seemed to show a high penetrance in mixed breed dogs with characteristic clinical signs such as dizziness, ataxia, weakness in the hind limbs (particularly in the morning hours), seizures, and even aggression and blindness reported in three of four evaluated dogs. The fourth patient was described in the EMR as having an episode of hindlimb lameness, which may be suggestive of SCA, but follow up examination/diagnostics were not available.

In contrast to the severe signs in the above disorders, while the forms of ichthyosis originally characterized in Golden Retrievers and American Bulldogs (32, 33) are not as life threatening, they were correlated with the anticipated clinical signs in all dogs genetically at risk evaluated. These dogs presented with flaky, scaly skin as puppies, with some owners reporting seeing flaking from birth and the majority of owners noticing ‘dry skin’ at the first puppy exam (approximately eight weeks). In many cases, these dogs did present with multiple dermatologic episodes throughout the timeframe evaluated, showing secondary infections, pyoderma, alopecia, and otitis along with generalized scaling.

## Discussion

With the present study we aimed to expand knowledge on the distribution, genetic prevalence, and clinical relevance of an unprecedented number of canine disease variants in a dog population of unparalleled size, focusing on previously identified polymorphisms that largely follow a Mendelian mode of inheritance. We further explored the link between genetic inbreeding levels, measured on an individual dog level, and both Mendelian disease risk.

In line with our previous investigation of 152 disease variants in around 100,000 dogs (7), we find that most tested disease variants are shared by both the mixed breed and the combined purebred population. The most notable effect of a 10-fold increase in study population size and the addition of nearly 100 tested variants to the screening panel is that the proportion of variants that are found in both the mixed and purebred group has more than doubled compared to our previous study (from 41% to 88%) while fewer variants remain exclusive to either subgroup. This observation is explained in part by our conscious decision to include several variants with inconclusive evidence of causality beyond the original discovery breed for genotyping in the present study, but it is also likely that a larger mixed breed sample captures a wider variety of breed backgrounds and disease alleles segregating in the population. The proportion of variants not encountered in the study sample remained comparable to our previous study at approximately 17%, despite the increased number of tested variants. The collective group of non-observed variants consisted of several X-linked disease variants (e.g., hemophilia A and B; Supplementary Table 1), and variants originally characterized as private to specific cases or family lines (e.g., Alexander disease in Labrador Retriever, osteochondromatosis in American Staffordshire Terrier and congenital eye malformations in Golden Retriever (34–36)), and we consequently confirm the absence of these variants from the broader canine population. Another notable observation compared to our prior canine disease variant screening studies is that with the increase of the number of tested variants to 250, it is now more common than not for a dog to carry at least one variant potentially influencing disease risk (57.2% of dogs). This estimate highlights how management of inherited disease variants in breeding programs of purebred populations requires thorough screening of known disease variants with a particular focus on the variants that are most relevant within each breed, combined with simultaneous consideration of genetic diversity and the viability of the gene pool in breeding selections.

We, in collaboration with other investigators, have previously shown that breeds with higher-than-average inbreeding levels required greater amounts of veterinary care on average (37), and that purebred dogs as a group are more likely than mixed breed dogs to be homozygous for a common recessive disease variant (7). In the present study, we have for the first time been able to evaluate the correlation between a molecular measure of genome-wide inbreeding and overall Mendelian disease load on an individual level. We most notably observed a relationship, albeit one with a small correlation coefficient, between genome-wide reduction in genetic diversity and an increased risk of homozygosity for multiple autosomal/recessive disease variants in dogs. This finding provides a direct measurement-based example of how increased genomic homozygosity, e.g. as a consequence of the mating of close relatives in a closed gene pool, could contribute to driving inbreeding depression through the accumulation of deleterious recessives. Notably, mixed breed dogs as a group are no exception to the observed correlation between diversity levels and disease variants. While it is theoretically likely that disease variants are maintained in the mixed breed population through random breeding with frequencies varying over time due to random genetic drift, matings between closely related individuals are known to occur in settings such as puppy mills. This explains the very broad spectrum of diversity levels observed in mixed breed dogs and emphasizes the utility of recessive disease testing also in this group in a veterinary clinical setting.

While we show that disease variants are collectively common in dogs, we also note that the majority of the studied specific disease variants have a frequency of <1% in the population, making it difficult for any individual veterinary clinician to recognize, and maintain proficiency with, the broad spectrum of characterized inherited disorders. This stresses the need for continuing medical education and veterinary specialists trained within the field of genetic counseling, while highlighting the value of broad diagnostic screening technologies. The most common disease variants observed in the present study expectedly included some known genetically widespread variants that are also frequently requested targets for genetic screening, such as DM, CEA, prcd-PRA, HUU, exercise-induced collapse (EIC), MDR1 (multidrug resistance 1) medication sensitivity, and von Willebrand’s disease type 1 (vWD 1 (13–16,38–40). Despite extensive genetic testing for these variants, challenges in determining and understanding their penetrance and expressivity in different breed backgrounds partially persist, as exemplified by phenotype studies of dogs genetically at risk of DM, CEA and cone-rod dystrophy (cord1-PRA/crd4) (41–43). It can also be particularly difficult to interpret what role in the regulation of disease onset is played by some variants originally published based on study of a subset of the canine population, or with limited functional evidence supporting causality. Within the scope of this study, we present selected clinical validation case studies that support the relevance of specific variants on diverse genetic ancestry backgrounds, while laying the foundation for future prospective reports addressing additional late onset conditions as our cohort of tested dogs ages. We show that some variants previously associated with bald thigh syndrome in sighthounds, dilated cardiomyopathy (DCM) in Doberman Pinschers, and Bulldog-type cystinuria are very common in the broader canine population and may represent non-causal markers in themselves. We caution against making conclusive health care or breeding related decisions based on genetic testing for such variants alone in dogs of mixed breed ancestry or in other pure breeds until further studies are conducted. Investigations of the impact of putative genetic risk factors in dogs representing a diverse genetic ancestry background are essential for establishing variant causality and broader clinical relevance of genetic testing. Among other findings, we provide evidence suggesting in particular that the specific variants previously associated with canine leukocyte adhesion deficiency type III, *ITGA10*-related chondrodysplasia, hypocatalasia, factor VII deficiency, cone-rod dystrophy 2, *KCNJ10*-related spinocerebellar ataxia, both *PNPLA1* and *NIPAL4*-related ichthyoses, focal non-epidermolytic palmoplantar keratoderma, hemophilia A, *FAM83G*-related hereditary footpad hyperkeratosis, *VPS11-*related neuroaxonal dystrophy, skeletal dysplasia 2 or trapped neutrophil syndrome have high or complete penetrance across breed ancestry backgrounds, which suggests that these variants always bear relevance when present in a dog. Taken together, the body of evidence we have collected contributes to establishing genetic counseling and veterinary care guidelines related to canine disease variants. Breeding decisions informed by observations collected across the broad dog population are critical both to avoid producing offspring at risk of a disease, and to ensure that dogs carrying a non-causal variant are not unnecessarily excluded from breeding programs.

The present study represents the largest cohort of dogs, and the most extensive set of Mendelian disease variants, examined in a single study to date. The study sample is broadly representative of the breed composition in the canine pet population due to ascertainment both via direct-to-consumer genetic testing and DNA sampling during primary veterinary care which was largely carried out without prior suspicion of patient susceptibility for a specific genetic disease. Moreover, the cohort includes representation of more than 250 different breeds or breed varieties, and more than 150 countries or geographic territories, and we are therefore confident that the generated dataset is a valuable resource for the community that enables breed health committees and researchers to examine variant frequencies by breed and region in more detail. Large cohorts with both genotyping data and detailed medical history available are an invaluable resource for studying disease etiology and epidemiology, and while we note that the group of dogs with electronic health records accessible to us was of a young median age (1.17 years at the last recorded clinic visit), we lay a solid foundation for a prospective study. When interpreting the clinical validation outcomes and estimates of variant penetrance, it should nonetheless be noted that some genetically affected dogs may only go on to develop signs of disease later in life or might have been lacking the specific laboratory examination required for a definitive diagnosis. While our cohort provides the best available insight into the Mendelian disease heritage of the general dog population, we specifically acknowledge that our definition of a “purebred” (a genetically uniform group of dogs with a consistent, predictable conformational and behavioral type) also included dogs that did not have a documented pedigree as defined by a formal breed registry. Moreover, as for any diagnostic test and even with the strictest quality control measures, some potential for individual spurious false positive test results exists without follow up with a secondary genotyping technology such as sequencing across the whole cohort. We have focused on presenting only the most relevant disease variant findings (>1% frequency and more than one carrier of the same variant observed) in additional breeds for these reasons.

In conclusion, we report that it is relatively common for a dog to be at least heterozygous for one of the many known putative canine disease variants, while the majority of individual variants are very rare in the overall population. While many variants are enriched in specific breeds and absent from others, we conclude that health issues caused by Mendelian disease variants are broadly shared across both mixed and purebred populations, as we find that several variants cause similar signs of disease independently of breed ancestry. Understanding of the clinical relevance of variants that are already targets of commercial genetic tests represents an important advance towards creating a variant significance classification system for dogs and contributes to the enablement of precision medicine. As more disease variants are identified and screened in the future, it is likely to become inevitably clear that all dogs carry a number of deleterious recessive alleles. Maintenance of population diversity through sustainable breeding selections, outcrossing programs, and avoiding the overuse of specific popular sires represents a simple remedy that keeps breeds viable by counteracting the augmentation of recessive disease alleles.

## Materials and Methods

### Study sample

A total of 1,054,293 dogs (811,628 mixed breed and 242,665 purebred dogs) were genotyped as a part of this study. All DNA samples were owner-provided non-invasively collected cheek swab samples voluntarily submitted for commercial genetic testing (Wisdom Panel™, MyDogDNA™, and Optimal Selection™ Canine genetic screening products) by Wisdom Panel (Portland, OR, USA) between November, 2019 and August, 2021. The study cohort represented a subset of the total more than three million dogs of varying ancestry genetically tested at Wisdom Panel, genotyped on a cross-compatible microarray technology platform. All owners provided consent for the use of their dog’s DNA in research. The country of origin of each dog was defined as the country where the sample was submitted for genetic analysis, unless specific information stating otherwise was reported by the owner. A total of 160 countries or autonomous regions were represented in the dataset (96 regions with >5 dogs). The vast majority of dogs (93.9%) were from the United States, with other notable subgroups being dogs from the United Kingdom (2.5%), Germany (1.2%), France (0.5%), Australia (0.3%), Finland (0.2%), and Canada (0.2%).

As one major motivation for clients to pursue Wisdom Panel™ genetic testing is gaining insight into their dog’s breed ancestry, the purebred status of a dog was either not known or considered prior to genotyping. For the purposes of this study, a dog was considered “purebred” if its genetic testing results indicated at least 7 of 8 great-grandparents being purebreds of the same breed. Notably, we did not strive to use a definition of “purebred dog” synonymous with the term “pedigreed dog” in terms of eligibility for registration with a recognized kennel club or breed registry. The purebred cohort (Supplementary Table 2) consisted of 263 different breeds or breed varieties (218 breeds represented by >5 dogs) and 34 archived samples from wild canids (Gray wolves, Dingos and Coyotes). The composition of the cohort was determined by the breed types routinely submitted for testing with Wisdom Panel™. Breeds contributing more than 2% of individuals of the overall study sample were: American Staffordshire Terrier (17.6%), Labrador Retriever (6.9%), German Shepherd Dog (6.4%), French Bulldog (5.4%), Golden Retriever (5.3%), Siberian Husky (3.7%), Yorkshire Terrier (3.4%), Shih Tzu (3.1%), Border Collie (2.8%), Pomeranian (2.2%), Beagle (2.2%), Pug (2.1%), Chihuahua (2.2%) and Standard Bulldog (2.0%).

### Genotyping

Genotyping of 250 disease variants and 1877 genome-wide representative, and cross-microarray platform version available, single nucleotide polymorphism (SNP) markers used for genetic heterozygosity evaluation was carried out according to manufacturer-recommended standard protocols on a custom-designed Illumina Infinium XT microarray (Illumina, Inc., San Diego, CA, USA). Protocols for microarray design, validation, and data quality control have previously been described in detail (6,7,44). All genotyping data for samples with a genotyping call rate <97% was discarded in the scope of the present study. Mendelian variants were selected for analysis based on pre-existing evidence implicating the variants in the etiology of canine inherited disorders, either as summarized in the Online Mendelian Inheritance in Animals (OMIA) database or based on a literature review. A small number of more recently characterized genetic variants (N = 9; Supplementary Table 1) were only available on a later updated version of the custom microarray, and they were consequently genotyped in a subset of dogs (N = 46,062).

### Statistical analyses

All statistical modeling was carried out in Minitab® version 19.2020.1 statistical analysis software. Standard non-parametric tests were used for comparisons of heterozygosity levels between groups, as the variable was non-normally distributed. The Mann-Whitney U test was used for comparisons of two groups (e.g, mixed breed and purebred dogs), and the Spearman rank correlation test was used to evaluate the relationship between heterozygosity level and N disease variants carried in the heterozygous or homozygous state. For the correlation analysis, dogs carrying >5 variants were combined with dogs carrying 5 variants to obtain subgroup sizes of ≥20 dogs.

### Clinical data evaluation

Several approaches were taken to explore the relationship between genetic variants and their respective expected associated phenotypes. Data from electronic medical records (EMRs) were available for 458,433 of the genotyped dogs enrolled for Optimum Wellness Plans® at Banfield Pet Hospital® clinics during the years 2018 - 2021. The EMRs of dogs genetically at risk were assessed to uncover links between genotypes and relevant ailment phenotypes. Additional phenotype information was collected through submission of clinical documentation by dog owners or interviews with a Wisdom Panel veterinarian or geneticist. In addition, genetically affected dogs were recruited for clinicopathological investigations where possible.

## Acknowledgments

We deeply thank all pet owners and breeders who have supported research by submitting samples from their dogs for genetic testing and openly shared information about their dogs’ health, wellbeing, and conformation. We are grateful to the more than 1000 Banfield Pet Hospital clinicians who diligently and consistently recorded their observations in medical records. Without that collective effort this work would not have been possible. We also wish to thank Dr. Hannes Lohi and team members (University of Helsinki), Dr. Eva Furrow and team members (University of Minnesota), and Dr. Cathryn Mellersh and team members (University of Cambridge) for sharing their knowledge on specific variants assessed in this work. Our colleagues Dr. JoAnn Morrison, Susan Pearce-Kelling, Nathaniel Spofford, and Dr. Jason Huff are thanked for their excellent advice. Finally, we wish to extend a special thanks to all our research colleagues and peers, whose dedication to the discovery of genetic variants underlying canine disease has made studies like ours possible.

## Supplementary material

Supplementary Table 1. Allele and genotype frequencies, and prevalence ranks for 250 screened variants in the full study sample.

Supplementary Table 2. Study sample composition and descriptive heterozygosity statistics by breed.

Supplementary Table 3. Summary of disease allele frequencies across breeds.

Supplementary Table 4. Descriptive information and phenotype details of clinically evaluated dogs.

